# Histone Modification Metapeaks are Epigenetic Landmarks Predictive of Cell State

**DOI:** 10.64898/2026.03.31.715657

**Authors:** R. Matthew Tanner, Theodore J. Perkins

## Abstract

Histone modifications are a key component of the epigenetic state of a cell, and they vary widely across different cell and tissue types, conditions, and disease states. Indeed, the majority of the genome is enriched with one histone mark or another across the thousands of cellular conditions that have been studied to date. Here, we use the largest-to-date collection of histone modification ChIP-seq datasets to identify the most important sites of histone modifications genome-wide. Collected and uniformly reprocessed by the International Human Epigenome Consortium, this data includes 5339 datasets enriched at nearly one billion total peaks across 59 different major cell or tissue types and in healthy and disease conditions, for six different histone marks. We propose FindMetapeaks, a new approach to identifying histone mark metapeaks, which are genomic regions with enrichment of a mark across many samples. We show that many of these epigenetic metapeaks are strongly indicative of cell and tissue type, or are associated with other sample characteristics, and highlight key regulatory regions of the genome. However, we also show that many metapeaks contain redundant information, and that parsimonious subsets of metapeaks can be selected by machine learning to predict cell state. Our histone mark metapeak atlas provides a concise set of regions for interpreting the epigenome.

**Availability:** https://github.com/rmbioinfo83/FindMetapeaks/

## 1 Introduction

Post-translational modifications to histones, or histone marks, are a key part of the epigenetic state of the cell (Bártová et al, 2008; Gates et al, 2017; Talbert and Henikoff, 2021). Hundreds of distinct modifications have been identified, many of whose exact functions are still under investigation. Still, a number of key marks are relatively well understood and have important functional implications for the cell. For instance, histone 3 lysine 27 acetylation (H3K27ac) marking of either promoter or enhancer regions is associated with gene activation (Creyghton et al, 2010). Conversely, trimethylation (H3K27me3) of promoters or gene bodies is associated with poised or repressed genes (Young et al, 2011; Wiles and Selker, 2017). Similar to how different cell types and disease conditions are reflected in distinct patterns of gene expression (Ross et al, 2000; McKenzie et al, 2018), so too do distinct patterns of histone marks at different genes associate with cell type or disease state (Heintzman et al, 2009; Greer and Shi, 2012).

The most common method for genome-wide mapping of histone modifications is ChIP-seq (Robertson et al, 2007; Park, 2009), which works by cross-linking DNA to the histones and then immunoprecipitating that DNA, using an antibody that specifically recognizes one type of histone modification. Successors to ChIP-seq include ChIP-exo (Rhee and Pugh, 2011), CUT&RUN (Skene and Henikoff, 2017), and CUT&Tag (Kaya-Okur et al, 2019), which aim for greater precision or signal-to-noise ratio. Regardless of the technology, all approaches result in DNA fragments from the vicinity of the histone modification of interest. These fragments are sequenced and mapped to the genome. Typically, regions of the high fragment density or enrichment, possibly compared to some control experiment, are identified by peak calling (Zhang et al, 2008; Thomas et al, 2017). In modern ChIP-seq data for histone marks, one typically finds tens or hundreds of thousands of enriched peaks. What one does with those peaks depends on the purpose of the study, and may include: comparison to peaks for other histone marks or transcription factors; comparison to other genomic features like genes, promoters, enhancers, or repeat elements; analysis for motif enrichment or conservation; association to nearby genes, to understand possible regulatory targets, etc. (Bernstein et al, 2005; Lee and Mahadevan, 2009; Ngo et al, 2019).

Numerous research projects have studied one or a few histone marks in one or a small set of samples. However, there have been several large-scale efforts to systematically produce histone mark data for many samples, and for a diversity of marks. The Roadmap Epigenomics Consortium generated a collection of “reference epigenomes” which included multiple histone marks as well as DNA methylation and other data, for 111 distinct cell or tissue samples (Bernstein et al, 2010; Kundaje et al, 2015). The Blueprint Epigenomics project studied the epigenetics of blood in great detail, generating histone mark and other data for nearly 500 individual human donors and seven cell lines (Adams et al, 2012). The ENCODE consortium has, as of this writing, generated 3694 histone-mark ChIP-seq datasets, mostly in human, and spanning a diversity of marks and tissue types, although as yet there is no integrated analysis of all that data (Sloan et al, 2016). Recently, the International Human Epigenome Consortium (IHEC) (Stunnenberg et al, 2016), drawing on Roadmap, Blueprint, and ENCODE datasets as well as significant new data generation, has put together and uniformly (re)processed a collection of 5339 ChIP-seq experiments covering six major histone marks across 1698 unique human samples.

With up to hundreds of thousands of peaks per dataset, and thousands of datasets, enormous collections such as these can contain hundreds of millions of regions of histone modification. The IHEC collection in particular contains nearly a billion, comprising 953,881,151 peaks. This presents a challenge for extracting biological meaning, but also an opportunity to identify the most “important” sites of epigenetic modification. What is important depends on one’s interests, but in general it should be sites that are consistently marked across a significant portion of the samples, and whose epigenetic status is statistically associated with different tissue types, or different disease states or prognoses, or the activity of key genes or pathways, etc. Here, we analyze the IHEC histone mark peaks using a novel approach to metapeaks identification, to produce a shorter list of candidate regions of greatest interest, and then further analyze those regions by statistical and machine learning methods for associations to covariates of interest.

The term “metapeaks” has been used in numerous studies where multiple ChIP-seq or similar datasets are being analyzed. Sometimes, metapeak analysis has meant plotting the average ChIP-seq signal, or a signal heatmap, across all peaks in a set or relative to some other genomic loci (González et al, 2015; Keller et al, 2021; Savadel et al, 2021). This can provide useful visualizations and insights, but is too cumbersome when one has hundreds or thousands of datasets to summarize. In other cases, metapeak analysis has meant taking the union of peaks in different datasets (Eaton et al, 2011). Conversely, the ENCODE project pipeline utilizes Irreproducible Discovery Rate analysis (Hitz et al, 2023), which combines replicate ChIP-seq peak sets by intersection, and then further narrows that list by ensuring that overlapping peaks in different replicates have similar statistical significance. Understandably, union and intersection strategies are respectively too loose and too stringent when dealing with hundreds or thousands of datasets. Meuleman et al (2020) recently published a large collection of DNase-hypersensitivity datasets, reporting regions of chromatin accessibility for over 400 different tissue types. Their data analysis included a clustering procedure to group closely overlapping peaks from different datasets, generating what could be called metapeaks of DNAse hypersensitivity. Their analysis used relatively stringent criteria for allowing peaks to cluster together, reducing the 76 million total regions they found across all their datasets to 3.6 million common regions. Unpublished work by Kudron et al (2024) proposes a metapeak mechanism similar to what we developed and propose here. We compare their method with ours in more detail in the Discussion section, but both share the same core idea: that the peaks arising from many individual ChIP-seq datasets can be treated as if they were individual mapped DNA fragments, and peak calling can be applied again to them. That is, we can use existing peak calling algorithms to identify peaks of peaks—a natural notion of metapeaks. This basic method is a sensible intermediate to either intersection or union strategies, while producing a shorter list of broader regions than the approach of Meuleman et al (2020).

The IHEC data we analyze comprises 5339 samples spread across six histone marks: H3K27ac, H3K27me3, H3K36me3, H3K4me1, H3K4me3, and H3K9me3. All the data is from human primary samples or cell lines. The data includes samples both from healthy and diseased conditions, including many cancer samples, spread across numerous cell and tissue types, ages, and males and females. We apply our FindMetapeaks strategy to call peaks of peaks for each histone mark individually. These metapeak sets are vastly less complex than their source datasets, comprising roughly two to three orders of magnitude fewer genomic regions. Yet, each metapeak set retains properties that are expected for the mark, such as associations to gene bodies, promoters or enhancers. Moreover, we find that many metapeaks are statistically associated to specific cell or tissue types, and that this is true for all six marks—not just those, such as H3K27ac, that are already well known for cell type specificity. We also find that within certain tissue types, a comparison of healthy versus cancer samples identifies cancer-enriched metapeaks that are tissue specific. We find histone mark metapeaks associated with many well-studied and crucially important genes, such as P53 (Brady and Attardi, 2010), Yamanaka factor genes (Takahashi and Yamanaka, 2016), and master transcription factors for various cell types. We propose that these metapeaks may provide a core set of genomic regions that can be used to interpret future histone mark datasets, and provide a concise coordinate system for representing differences among those datasets.

## 2 Results

### 2.1 Metapeaks as “Peaks of Peaks”

In order to identify key genomic regions of interest across a large number of ChIP-seq or related datasets, we propose a new approach called FindMetapeaks (https://github.com/ rmbioinfo83/FindMetapeaks/). This approach is outlined in Figure 1, which also puts it into context with traditional ChIP-seq peak calling. In traditional peak-calling one compares the density of ChIP-seq reads along the genome with some control read density. Regions where the ChIP-seq reads are of significantly greater density than the control reads are labeled as peaks. There are numerous peak calling algorithms, with MACS2 being the most widely used at present (Zhang et al, 2008; Feng et al, 2012; Rashid et al, 2011; Shen et al, 2013; Meers et al, 2019). In our approach, metapeak calling begins with *N* different collections of peak sets, each generated by performing peak-calling on *N* different ChIP-seq datasets and their matched controls. In order to account for differences in the number of peaks called per dataset, and avoid any single dataset or subset of the data having disproportionate influence, we retain the top *P* peaks per dataset, as ordered by decreasing statistical significance. (This assumes the peak-caller outputs not just the peak regions, but some p-value or score that can be used for ranking; this is true of MACS2 and all other peak callers we know of that are in common use.) The top *P* peaks per sample are collected into a list of *N × P* total peaks, to which peak calling is applied again, identifying “peaks of peaks”, or metapeaks. In this case, there is no control set to which one compares, so it is necessary to use a peak caller that is capable of calling peaks without a reference control, such as MACS2. When no control is present, MACS2 compares the inputted regions to an imagined uniform distribution with the same number of regions, as well as to a locally-varying background based on the input regions. It calls peaks (in our case, metapeaks) wherever the input regions (in our case, peaks) are significantly denser than the uniform or locally-varying backgrounds. Once metapeaks have been calculated, many analyses are possible. In particular, for several analyses we re-represent the original samples in terms of a metapeak matrix. This matrix, of size number of samples (*N*) by number of metapeaks, can be either binary or integer, representing respectively whether or not each sample has a peak overlapping each metapeak, or representing how many peaks each sample has overlapping each metapeak. This matrix representation of all the ChIP-seq datasets is a key benefit of metapeak calling, as otherwise there is no obvious, concise, fixed-sized coordinate system in which to compare different samples—a key motivation behind the Meuleman analysis as well (Meuleman et al, 2020).

**Fig. 1.**
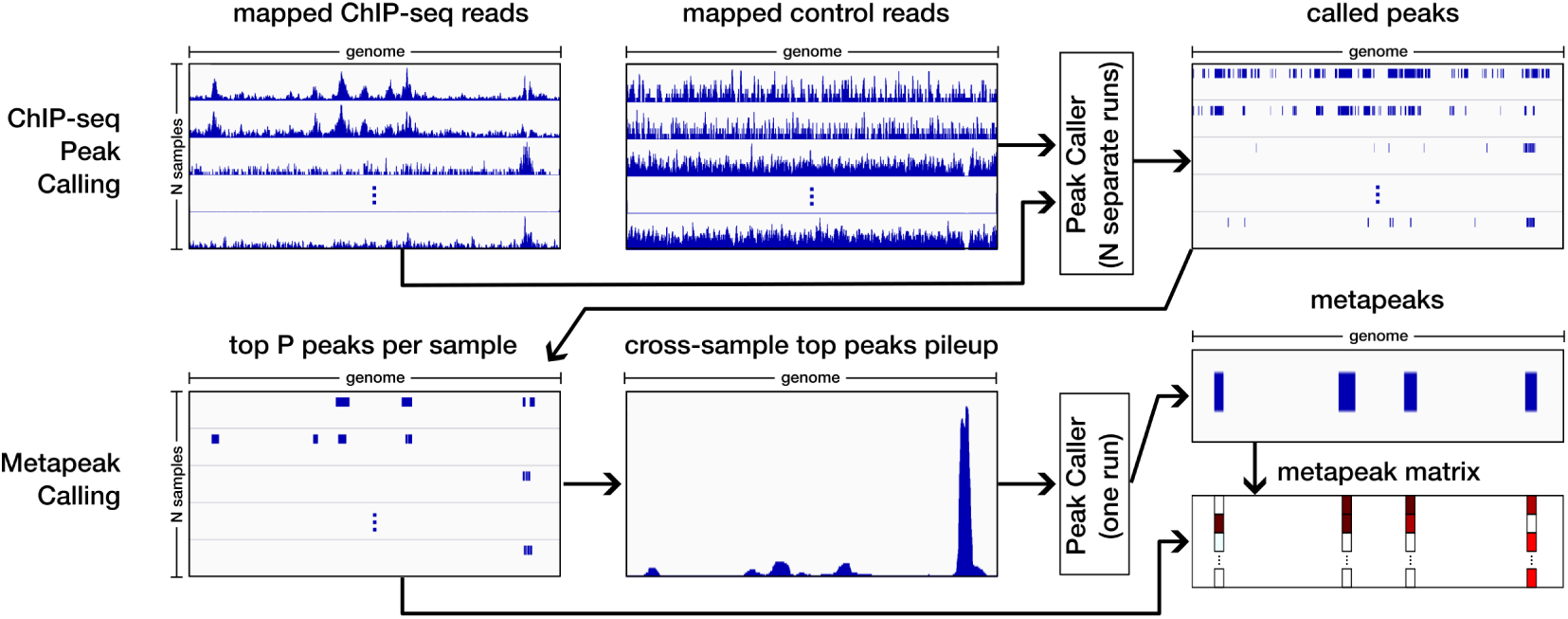
Outline of our FindMetapeaks strategy and its relationship to traditional single-dataset peak calling. Peak calling for ChIP-seq or related types of data compares densities of mapped reads in a ChIP-seq or “treatment” dataset versus a control dataset. Regions where ChIP-seq reads are significantly higher than control are termed peaks. Metapeak calling uses the peaks called from many individual ChIP-seq experiments, treating them as if they were mapped reads. In our FindMetapeaks approach, the top *P* most significant peaks from each of *N* ChIP-seq peak sets are selected, in order to correct for imbalance in numbers of peaks per dataset. The selected peaks are combined and subjected to peak calling without any control, and the “peaks of peaks” thus identified are the metapeaks. By determining if or how many top *P* peaks from each sample overlap with each metapeak, we produce a metapeak matrix, which represents the samples in metapeak space.

### 2.2 The International Human Epigenome Consortium Histone Marks Compendium

We applied FindMetapeaks to identify key regions of epigenetic modification in the human genome by applying it to six different collections of ChIP-seq datasets prepared by the International Human Epigenome Consortium (IHEC) (Supplementary File 1). These datasets span a number of cell and tissue types, ages, health and disease conditions, and comprise six major histone marks: H3K27ac, known as an activation-associated mark at gene promoters and enhancers (n=1565 datasets); H3K27me3, associated with gene repression (n=675); H3K4me1 and H3K4me3, which are associated with weak and strong gene activation at promoters and/or enhancers (n=963 and 799 respectively), H3K36me3, known for marking actively transcribed gene bodies (n=695); and H3K9me3, associated with repression in repetitive regions of the genome (n=642) (Talbert and Henikoff, 2021).

The IHEC metadata group divided the samples into 59 cell or tissue types. Some are very broad, such as “brain” (n=473), and are expected to harbor many individual cell types in unknown proportions. Others are far more specific, such as “T cell” (n=657), “neutrophil” (n=492), or “kidney epithelial cell” (n=3). The division into cell and tissue types depended in part on how many samples of different types were available, but also the availability or quality of metadata originally deposited along with ChIP-seq data, and the feasibility of mapping experimenter-provided metadata into standard Uberon (Mungall et al, 2012) or other ontology terms (Raby et al, 2025). The metadata also included a coarse categorization of samples into healthy (3868 samples), those having some type of cancer (1165), or those with other diseases (306).

For each ChIP-seq and control dataset, IHEC mapped all reads and called peaks using MACS2. Large variability in the number of peaks per sample was observed, ranging from just over 10k to 500k. Different numbers of peaks per dataset can result from any number of technical features rather than biological ones, including something as simple as sequencing depth. Therefore, to minimize biasing our analysis towards datasets with more peaks, we restricted our FindMetapeaks analysis to the top *P* = 10,000 peaks per sample, ranked by decreasing MACS2 peak scores (which are proportional to the negative log p-values). In cases where multiple peaks in a dataset were tied for having the 10,000th-best p-value, we took all such tied peaks, along with everything more statistically significant. This ensured that each sample was roughly equally represented. We used MACS2 for the metapeak calling, in no-control mode, and with other parameters at their defaults. Peaks were passed to MACS2 in the form of bedGraph files, and the MACS2 “bdgpeakcall” function was called.

### 2.3 Examples of Histone Mark Metapeaks

We used FindMetapeaks to call metapeaks separately for each histone mark, with the results available in Supplementary File 2. Because the FindMetapeaks notion of metapeaks for histone marks is a new one, we provide several intuition-building examples in Figure 2, before studying them more comprehensively in the remainder of the paper. Three different genomic regions are displayed, and peak pileups along with called metapeaks are shown for all six histone marks.

**Fig. 2.**
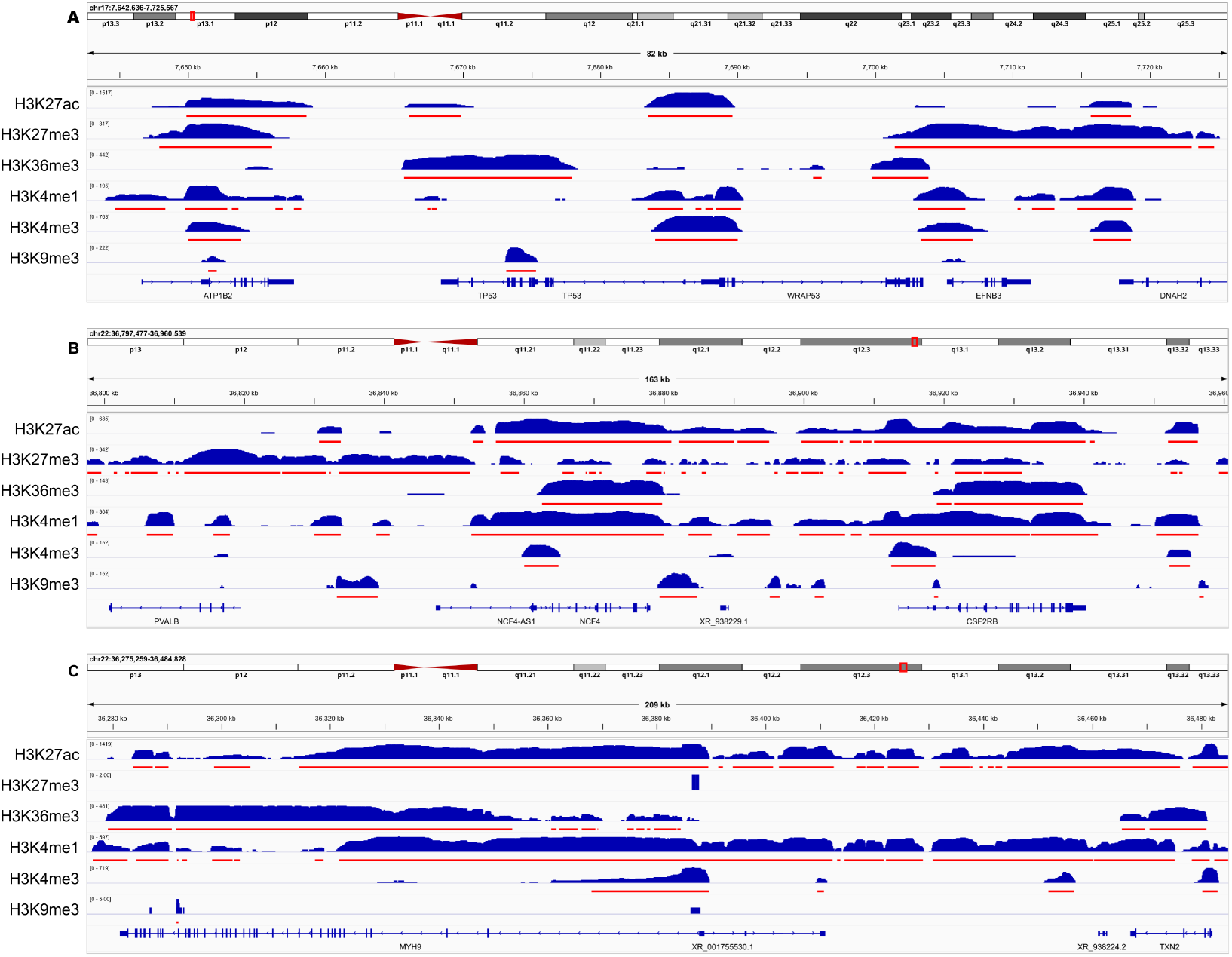
Example metapeaks for the histone marks H3K27ac, H3K27me3, H3K36me3, H3K4me1, H3K4me3, and H3K9me3 show a variety of broad and narrow peaks with differing epi-allele frequencies. The blue tracks are pileups of peaks from hundreds or thousands of ChIP-seq datasets, shown on a log axis and autoscaled to fill the vertical space, on which peak calling was applied again to identify metapeaks. The red bars show the identified metapeaks. Three different regions of the genome are shown, highlighting the vicinities of the genes (A) P53, (B) NCF4, and (C) MYH9.

Figure 2A shows metapeaks in the vicinity of Tumor Protein P53, a gene that is well known for its key roles in regulating the cell cycle, DNA repair, and apoptosis, and a gene that is especially important in cancer (Olivier et al, 2010). The panel shows substantial metapeaks for the activation-associated marks H3K27ac and H3K4me3 at the transcription start site (TSS) of the gene. There is also a metapeak for H3K36me3 on the gene body, which again is typically associated with active transcription. That metapeak falls especially on the exon-dense 3’ half of the gene, consistent with previous observations that H3K36me3 especially marks exons Kolasinska-Zwierz et al (2009). There is no metapeak for H3K27me3 on P53, suggesting that the epigenetic regulation of the gene is primarily through selective or condition-specific activation, rather than repression.

Figure 2B shows metapeaks in the vicinity of the gene Neutrophil Cytosolic Factor 4 (NCF4), which is expressed in a variety of tissues but especially the bone marrow and blood lineage cells, particularly neutrophils (Ryan et al, 2014; Pontén et al, 2008). (We return to the issue of tissue-specificity of metapeaks, including this neutrophil-enriched example, in Figure 4.) This gene is rich with metapeaks, including an H3K27ac metapeak that spans the entire gene, and an H3K36me3 metapeak that slightly more specifically covers the gene body, and an H3K4me3 metapeak centered on the TSS. These all point to strong epigenetic activation of the gene in various conditions. However, there are also some smaller H3K27me3 metapeaks on the gene, which might point to epigenetic repression in some contexts—although these may relate to overlapping gene NCF4-AS1 instead. Further to the left of Figure 2B we see the Parvalbumin gene (PVALB), which is involved in muscle relaxation and neurodevelopmental disorders, among other things (Rall, 1996; Janickova and Schwaller, 2020). This gene shows a very prominent H3K27me3 metapeak at its TSS, suggesting its epigenetic regulation is primarily repressive. (That peak would look even more prominent if the pileups were not shown on a log scale, which we have done to ensure visibility of the evidence for the smaller metapeaks.)

Figure 2C focuses on the Myosin Heavy Chain 9 gene (MYH9), with roles in cell division and motility, and associations to a number of genetic illnesses (Pecci et al, 2018). Like the genes in previous panels, it has H3K27ac, H3K4me1, and H3K4me3 metapeaks (the last particularly focused on the TSS) and H3K36me3 metapeaks on its exons. There are no metapeaks for H3K27me3 or H3K9me3 on this gene. This region of the genome shows heavy marking by H3K27ac and H3K4me1 in the intergenic region on the right side of the panel, suggesting possible sites of regulatory elements. Indeed, the GeneHancer database (Fishilevich et al, 2017) indicates several enhancers in the region, supporting this hypothesis.

Although we will shortly become more comprehensive in our analysis, the examples in these regions demonstrate several general characteristics of metapeaks. First, they can vary widely in size. Some here are smaller than 2,000 base pairs while others are larger than 100,000 base pairs. There is also large variation in the number of individual peaks contributing to each metapeak, or relatedly, their “epi-allele frequency”—the fraction of samples that contribute a peak to a metapeak region. Although the epi-allele frequency cannot be easily read in the genome browser view, taller metapeaks generally contain more peaks from a larger fraction of the datasets. As with the peaks from which they are built, the metapeaks also have characteristic regions of the genome where they can be found, such as on the TSS, gene body, enhancer regions, etc.

Similar to peak-calling on individual samples, one might dispute whether certain of these metapeaks should be called or not. For instance, in Figure 2A the H3K36me3 track shows a small metapeak in the middle of the WD Repeat Containing Antisense to P53 gene (WRAP53). This one barely passes statistical significance and may be spurious; but on the other hand it might be biologically significant for some very specific subset of the samples, and therefore valid. As another example, the vicinity of the TSS of P53 has four metapeaks for H3K4me1, each reflecting one visible peak in the H3K4me1 peaks pileup. But, should these really be four separate metapeaks, or three, or one? It is difficult to know without further investigation. The other panels too show examples of small peaks that are called and slightly smaller ones that are not, or broader “domains” that are broken into multiple metapeaks. It would be possible to manipulate the significance threshold in MACS2 to include more, or fewer, of these regions among the metapeak calls, or to perform “broad” peak calling or otherwise stitch regions together. For our analysis, however, we will stick with the metapeaks called by MACS2 using default settings.

### 2.4 Metapeaks for All Six Histone Marks

Metapeaks summarize large collections of peaks into a much smaller number of key cross-sample regions. Figure 3A details, for each histone mark, the total number of peaks across datasets, and the number of metapeaks into which FindMetapeaks summarizes them. For instance, H3K4me3 has the fewest metapeaks (24044), while H3K4me1 has the most (148957). In general, the number of metapeaks is approximately two to three orders of magnitude less than the number of peaks that they summarize—a ratio roughly in line with the base peak calling itself, where a typical ChIP-seq dataset will have tens of millions of reads summarized into tens or hundreds of thousands of peaks. The number of metapeaks produced does not substantially correlate with the underlying total sample peaks they were called from (r=0.17, p=0.75 across the six histone marks). For example, H3K27ac has by far the most peaks, but three other marks had more metapeaks.

**Fig. 3.**
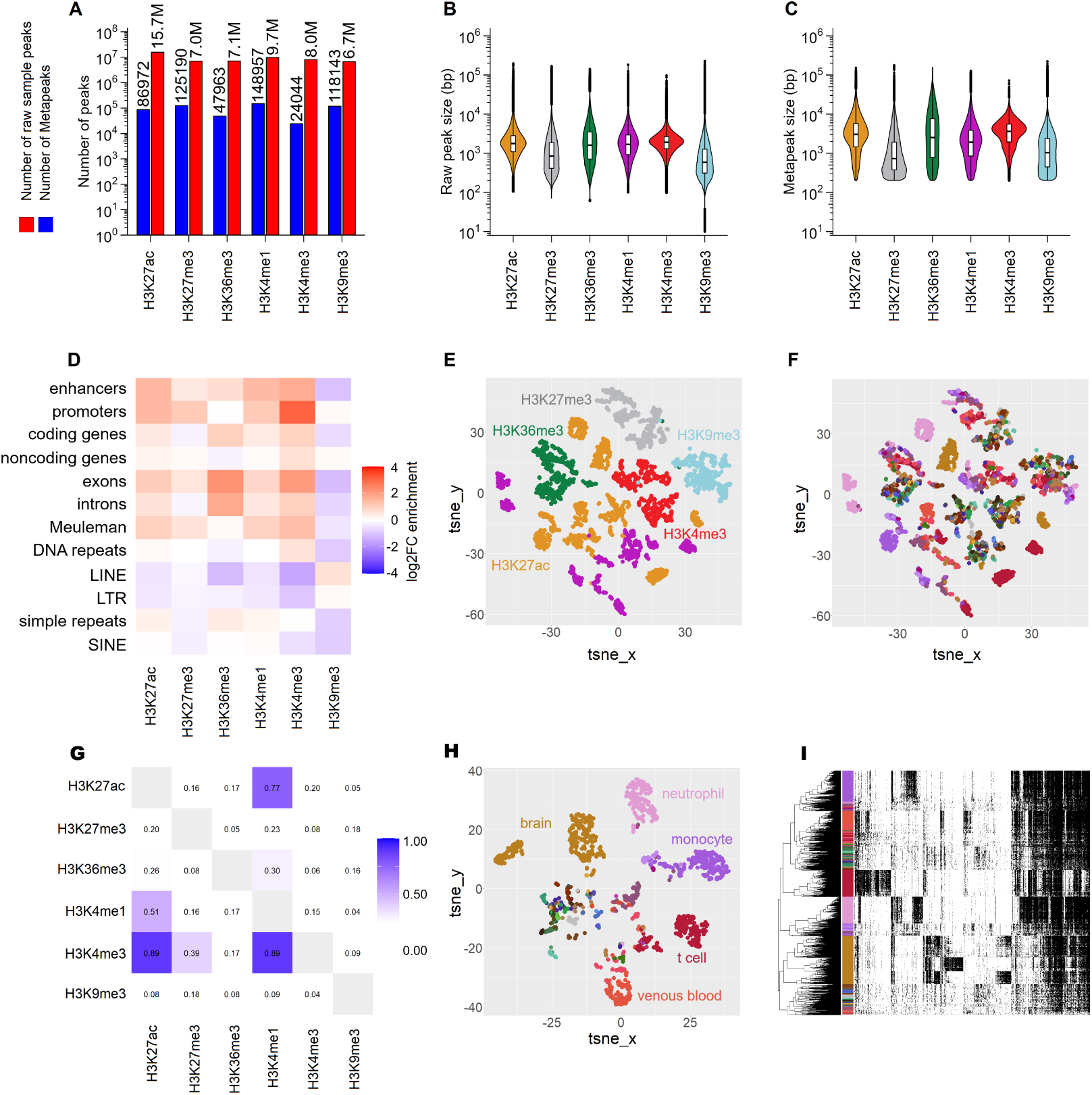
Statistical features of histone mark metapeaks mirror those of the source peaks, and display mark- and tissue-specific patterns. (A) Total numbers of peaks in the histone mark datasets versus the much smaller number of metapeaks into which they are summarized. (B,C) Violin plots showing size distribution of peaks (B) and metapeaks (C). (D) Relative enrichment of metapeaks over different types of genomic elements. (E,F) tSNE embedding of all metapeaks, colored by type of histone mark (E) or cell/tissue type (F). (G) Overlap analysis of metapeaks for different histone marks. (H) tSNE embedding of the H3K27ac metapeaks, showing clustering by cell/tissue type. (I) Binary heatmap of the H3K27ac metapeak matrix, showing cell/tissue-specific clusters of metapeaks.

Naturally, metapeaks are larger than the peaks from which they are built, but not vastly larger. Figure 3B,C show the distributions of peak sizes and metapeak sizes for the six different histone marks. Metapeaks are, on average, approximately 50% larger than peaks. For the activation-associated histone marks (H3K27ac, H3K36me3, H3K4me1, and H3K4me3), peaks are on average between 1-2kbp, while for the repression-associated marks (H3K27me3 and H3K9me3), peaks are below 1kbp average size. We note, however, that a small number of individual peaks are quite large, exceeding 100kbp. Metapeaks for activation-associated histone marks average 2-5kbp, while metapeaks for the repression-associated marks average less than 2kbp.

While the numbers and sizes tell us something about metapeaks, a key question is whether they contain meaningful biological information. In particular, one would hope that metapeaks would retain some key characteristics of the peaks themselves. As a first step in this direction, we characterized what types of genomic features metapeaks overlap, expressed as a fold change relative to shuffled regions (Figure 3D). Genomic features included enhancers as defined by GeneHancer (Fishilevich et al, 2017), open chromatin regions as defined by Meuleman et al. (Meuleman et al, 2020), gene promoters, bodies, introns, and exons according to GENCODE v29 (Frankish et al, 2021), and various types of repeat elements (Navarro Gonzalez et al, 2021). Enrichment fold change was calculated by comparing the number of metapeak regions overlapping feature regions compared to the number of shuffled metapeak regions overlapping those same feature regions. Generally, all marks except H3K9me3 are enriched on genes, promoters, enhancers, and open chromatin regions, and depleted in most types of repetitive DNA regions. The strongest enrichment observed was for H3K4me3 metapeaks on promoters, at roughly 16 times the level seen for shuffled regions. This is a sensible result given that H3K4me3 is known to mark active promoters (Wang et al, 2023). H3K27me3 metapeaks are also enriched at promoters, likely reflecting epigenetically silenced or bivalent genes (Blanco et al, 2020). H3K36me3 shows a strong enrichment for exons within genes, a quantitative result which substantiates the qualitative observation we made for H3K36me3 in Figure 2. H3K36me3 metapeaks are enriched for coding genes but not non-coding genes, which are slightly depleted.

H3K9me3 is the most distinct from the other histone marks. It is known to repress lineage-specific genes and to mark repeat-rich regions within the genome, including silencing retrotransposons (Zhang et al, 2024). We found some enrichment for H3K9me3 metapeaks on long terminal repeats, but in all other types of regions it was neutral or depleted compared with shuffled regions. Potentially, difficulties in mapping reads and calling peaks in repeat regions also affect our called metapeaks.

We also overlapped the metapeaks of each mark with those from every other mark. Figure 3G reports the fraction of the metapeaks for the mark named in the row that overlap any metapeak for the mark named in the column. H3K27ac, H3K4me1, and H3K4me3 by far overlap the most, consistent with their positioning at promoters and enhancers. H3K36me3 metapeaks also overlap those three marks somewhat, but to a lesser degree, consistent with their focus on gene bodies. Once again, H3K9me3 stands out as the most different histone mark.

The FindMetapeaks pipeline generates a matrix representation of the samples that can either represent the number of peaks each sample contributes to a metapeak, or simply a binary indicator of whether or not a sample contributes any peaks to a metapeak. We found that the majority of metapeaks in our analyses had either zero or one peaks from each sample, so we chose to use the binary metapeak occupancy matrices for all subsequent analyses. To represent all 5339 datasets in a common coordinate system, we used the bedtools merge function to combine the metapeak regions of all six histone marks into one master list of merged metapeak regions, and re-counted sample peaks into a master metapeak matrix. We generated a tSNE plot from this data, color coding each sample based on their histone marks (Figure 3E). The datasets cluster best by histone mark across the merged metapeak regions, rather than by cell type (Figure 3F). However, some clusters of T-cells, neutrophils, monocytes, venous blood, and brain are clearly visible (Figure 3F). These are the cell/tissue types with the largest number of samples overall. Pairwise correlation analysis confirms that histone mark is the most statistically significant influence on sample position in the embedding, followed by cell/tissue type (Figure S1).

To better visualize cell/tissue clustering, we chose to focus on only samples with the H3K27ac histone mark. A tSNE embedding based on the binary metapeak matrix for H3K27ac (Figure 3H) shows definite clustering of samples by cell/tissue type. A hierarchically clustered heatmap of the metapeak matrix (Figure 3I) demonstrates clear clusters of metapeaks that are highly enriched in specific tissue types (indicated by the colors at the left of the panel). For instance, brain (brown), neutrophils (pink), monocytes (purple), T-cells (red) and venous blood (red-orange) all show relatively specific metapeaks. Interestingly, the brain samples show two distinct groups of enriched metapeaks, which turn out to correspond to cerebellum and cortex samples. The next section more carefully analyzes the metapeaks for cell/tissue type enrichment.

### 2.5 Ubiquitous and Tissue-Enriched Metapeaks

The metapeak matrices for each histone mark have a row for every sample and a column for every metapeak. However, if samples can be categorized into discrete types (e.g. cell/tissue type, disease vs healthy, etc.) then we can produce a summary matrix of epi-allele frequencies (EAFs) by computing the fraction of samples of each type that have a peak in each metapeak. Equivalently, the EAF matrix is obtained by averaging the rows of the binary metapeak matrix within each sample type, producing a matrix indexed by sample types on rows and metapeaks on columns. For each histone mark, we computed EAF matrices based on cell/tissue type, restricting attention to 19 types with at least 10 samples in at least one of the histone marks. Figure 4A-F shows those matrices, restricted to a subset of metapeaks (columns) limited by the resolution of the image. The metapeak columns are hierarchically clustered based on Euclidean distance, while rows are cell/tissue types indicated by color, with full identities spelled out explicitly in Figure 4J. In the heatmap, colors from white to black indicate EAF values ranging from 0 to 1. Pink values denote tissues that did not have the requisite 10 samples for estimating EAF.

**Fig. 4.**
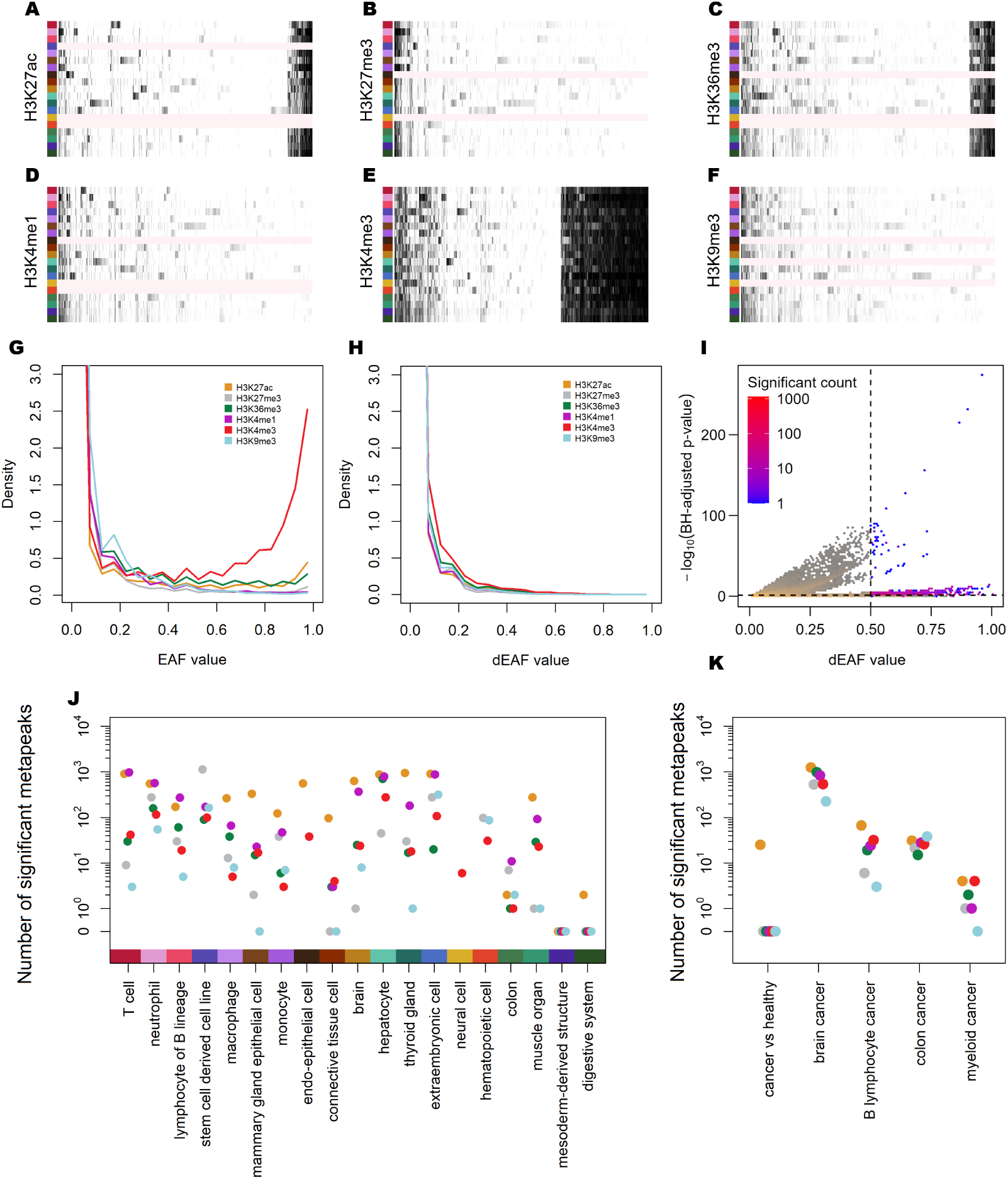
Epi-Allele Frequencies Reveal Ubiquitous and Tissue-Enriched Metapeaks. (A-F) Greyscale clustered heatmaps of metapeak epi-allele frequencies (EAFs) for each histone mark, for each cell/tissue type listed in Figure 4J. Cell/tissue types with insufficient sample size (*<* 10 samples) have no data shown, denoted by the light pink coloring. (G) Density plot of EAF values for each histone mark across all cell/tissue types combined. (H) Density plot of delta-EAF (dEAF) values for each histone mark across all cell/tissue types combined. (I) Volcano density plot of dEAF values versus -log10(q-values). (J) Number of uniquely upregulated metapeaks for each cell/tissue type, for each histone mark (q-value *≤* 0.05 & dEAF *≥* 0.5). (K) Significantly upregulated metapeaks for cancer vs healthy comparisons, for each histone mark (q-value *≤* 0.05 & dEAF *≥* 0.5).

Three of the gene expression-associated marks, H3K27ac, H3K36me3, and H3K4me3, show a subset of metapeaks that are marked strongly across all tissues. These appear at the right end of each heatmap. H3K4me3 has the greatest percentage of metapeaks of this sort. However, that is also the mark with the fewest metapeaks overall (∼24k). These can also be seen in Figure 4G, where we show density plots of EAFs for each histone mark. For all marks, the EAF density is highest near zero, between 69.2% and 82.3% of metapeak-tissue pairs having EAF *<* 0.05, depending on the histone mark. But several marks, H3K4me3 especially, have a high density of metapeaks with EAF near one.

We define “ubiquitous” metapeaks as those with a mean EAF ≥ 0.9 across all tissue types for a given histone mark. This identifies 3864 ubiquitous H3K4me3 metapeaks, 953 for H3K27ac, 321 for H3K36me3, 161 for H3K27me3, 154 for H3K9me3, and 9 for H3K4me1. This distribution is biologically plausible, because H3K4me3 and H3K27ac are strongly associated with active promoters and regulatory regions, whereas H3K36me3 marks transcribed gene bodies; by contrast, H3K27me3 and H3K9me3 are repressive marks that are generally more cell type- or locus-restricted, and H3K4me1 is most often associated with enhancer contexts rather than uniformly active promoter elements (Barski et al, 2007; Ernst et al, 2011). The ubiquity of the active marks suggests “housekeeping” functions, and Genomic Regions Enrichment of Annotations Tool (GREAT) GO Biological Process analysis supported this interpretation (Gu and Hübschmann, 2023) (Supplementary File 3), although the level of specificity differed by mark. For the large ubiquitous H3K4me3 and H3K27ac sets, the enriched terms were dominated by broad cellular and transcription-linked categories, including cellular macromolecule metabolic process, nucleic acid metabolic process, RNA metabolic process, gene expression, and mRNA metabolic process, suggesting that these ubiquitous active-mark metapeaks preferentially map to genes involved in core cellular maintenance and transcriptional output. The ubiquitous H3K36me3 set showed a similar but somewhat sharper emphasis on transcription-associated functions, with enrichments such as RNA processing and macromolecular complex subunit organization, again consistent with constitutively transcribed loci (Benayoun et al, 2014; Ernst et al, 2011). In contrast, the much smaller ubiquitous H3K27me3 and H3K9me3 sets were enriched for more regulatory and developmental terms, consistent with prior work showing that Polycomb-associated repression often targets developmental regulators, and that H3K9me3 can mark specialized repressed domains including zinc-finger gene clusters and other heterochromatic loci (Lee et al, 2006; Blahnik et al, 2011; Severson et al, 2013). Meaningful analysis of the H3K4me1 ubiquitous set by GREAT is not feasible as it contains only nine peaks.

Each histone mark also has metapeaks that appear enriched in specific tissues. These are visible as dark/black stretches of columns in one row that are largely white/light in the other rows (Figure 4A-F). This is true not only for H3K27ac, which is well known for tissue specificity, but for other marks as well. We posit two criteria to identify tissue-specific metapeaks. The first involves delta-EAF (dEAF), which we define as the EAF in a tissue minus the highest EAF observed among all remaining tissues for that same metapeak. This is meant to capture an “effect size” or extent to which one tissue has an EAF significantly higher than all the others. By construction, only one tissue can have a positive dEAF for a given metapeak—the tissue with maximum EAF, if there is a unique one. Figure 4H shows that the dEAF distributions are much more compressed toward zero than the EAF distributions, with density dropping rapidly by about dEAF 0.5 for all marks. Therefore, the first of our two criteria for tissue-specific enrichment is dEAF ≥ 0.5. However, this notion does not capture the confidence we have in the EAF or dEAF values. For example, suppose a tissue has exactly 10 samples with an EAF of 0.6 and dEAF of 0.5. Those EAF and dEAF values might change substantially if we took 10 new samples. But we would be more confident in the same statistics for a tissue with 100 samples. Therefore, our second test for tissue enrichment involves a Fisher’s exact test comparing the candidate-tissue’s EAF against all remaining tissues combined. This captures a notion of confidence in the observed proportions. The Fisher’s test is followed by a single Benjamini-Hochberg correction across all tests from all histone marks, and we require both q *<* 0.05 and dEAF ≥ 0.5 to assert enrichment.

The density-volcano plot in Figure 4I displays q-values and dEAFs for all candidate tissue-metapeaks pairs, with those passing both criteria highlighted in a blue-to-red gradient indicating how many pairs have those values. Two distinct significance regimes are evident. Most tissues with modest sample numbers produce associations with -log10(q-values) below 10, whereas tissues with much larger cohorts, such as T-cells, brain, and neutrophils, generate a second regime extending to far higher significance. For instance, several T-cell associations exceed -log10(q-value) of 200. This separation is expected from the marked differences in statistical power across tissues, while the dual-density rendering helps distinguish the full cloud of tested associations from the subset that passes both statistical and effect-size thresholds.

Across all six histone marks, we identified 15,764 significant metapeak-tissue associations, comprising 6645 for H3K27ac, 4462 for H3K4me1, 1969 for H3K27me3, 1198 for H3K36me3, 831 for H3K4me3, and 659 for H3K9me3 (Supplementary File 2). The number of associations varies widely by tissue (Figure 4J). For instance, there are: 912 T-cell-specific H3K27ac metapeaks and 942 for thyroid gland; 1,140 H3K27me3 metapeaks for stem cell-derived cell lines; and 703 H3K36me3 metapeaks for hepatocytes. At the opposite extreme, digestive system yielded only 2 unique H3K27ac metapeaks and none for the other marks, while mesoderm-derived structure yielded none across all six marks, foreshadowing the difficulty of resolving these tissue labels in the machine learning analyses in the following section.

Many genes nearby to metapeaks with tissue-specific enrichment are biologically plausible. For example, brain-associated regions recurrently mapped near neuronal genes such as Contactin-2 (CNTN2) (Bizzoca et al, 2003), Neurofascin (NFASC) (Buttermore et al, 2012), and Neurotrimin (NTM) (Struyk et al, 1995). T-cell-specific metapeaks were near canonical lymphoid genes such as Interleukin 7 Receptor (IL7R) (Akashi et al, 1998) and IL-2-inducible T-cell kinase (ITK) (Lucas et al, 2002). Neutrophil-linked metapeaks included C-X-C Motif Chemokine Receptor 2 (CXCR2) (Goncalves and Appelberg, 2002) and Bactericidal Permeability Increasing Protein (BPI) (Weiss and Olsson, 1987). Thyroid examples were particularly canonical, including thyroglobulin (TG) (Chambard et al, 1987) and Thyroid Stimulating Hormone Receptor (TSHR) (Marians et al, 2002). GREAT analysis of the tissue-enriched metapeaks from the four activating marks further supports this interpretation (Supplementary File 4). For instance, the top four hits for brain-linked metapeaks are: nervous system development (hyper-BH q-value *<* 10*^−^*^16^), neurogenesis (q *<* 10*^−^*^14^), generation of neurons (q *<* 10*^−^*^12^), and neuron differentiation (q *<* 10*^−^*^11^). Taken together, these examples and systematic GREAT analyses support the interpretation that the tissue-enriched metapeaks we are recovering constitute recognizable tissue-specific programs rather than arbitrary statistical outliers.

### 2.6 Cancer-Enriched Metapeaks

Of all the ChIP-seq peak sets in the IHEC data release, 1165 are annotated as coming from some kind of cancer sample, either a cancer cell line or a patient tissue/blood sample. An additional 3868 samples are marked as healthy. To see if there are any cancer-associated metapeaks, we first compared the group of all cancer samples against the group of all healthy samples, using the same criteria as described in the previous section—requiring dEAF ≥ 0.5 and Fisher’s exact test q-value ≤ 0.05. This pan-tissue comparison yielded only a few significant metapeak regions and these were all for H3K27ac (Figure 4K).

Despite their paucity, these findings are potentially meaningful. For instance, we found a cancer-enriched Paired Box 5 (Pax5) metapeak, a gene with a normal role in B-cell development, but whose aberrant expression is noted in B-cell and other cancers (O’Brien et al, 2011). We also found a metapeak at miR-155, which can play a role in numerous cancer types via mechanisms such as suppressing tumor suppressors, promoting the EMT, and promoting chronic inflammation (Tili et al, 2009; Mattiske et al, 2012; Bayraktar and Van Roosbroeck, 2018; Wu et al, 2023). There is a cancer-enriched H3K27ac metapeak at Musashi RNA Binding Protein 2 (MSI2), which has roles in a diversity of cancers (Kudinov et al, 2016; Guo et al, 2017; Li et al, 2020). Other cancer associated genes with metapeaks include Spi-B Transcription Factor (SPIB), Immunoglobulin Heavy Constant Mu (IHGM), and SRY-Box Transcription Factor 12 (SOX12).

Thinking that the diversity of tissues might be obscuring some findings, we also performed a cancer versus healthy analysis for four tissues that had at least 10 healthy and 10 cancer samples: brain, B-lymphocytes, colon, and myeloid cells. These analyses found additional cancer-enriched metapeaks, particularly in brain. One example is Lymphoid Enhancer Binding Factor 1 (LEF1), which, despite its name, is implicated in a broad array of cancers and promotes EMT (Li et al, 2006, 2009; Gao et al, 2014; Santiago et al, 2017). The cancer-enriched metapeaks are denoted in Supplementary File 2.

### 2.7 Machine learning combines biologically-relevant metapeaks for robust cell/tissue-type prediction

In Section 2.5 we showed that many metapeaks are enriched in certain tissues. However, this does not immediately imply that the tissue type of a sample can be determined definitively from its row in the metapeak matrix—i.e. from the list of metapeaks to which it contributes a peak. Even for the most strongly-enriched metapeaks, there are typically some peaks contributed from other tissue samples. Furthermore, our enrichment analysis does not speak to whether metapeaks provide the same redundant information, or if combining information across metapeaks might lead to greater discriminatory power, and which/how many metapeaks produce the best discrimination.

As a simple test of these questions, for each histone mark we trained L1-penalized multi-class logistic-regression classifiers to predict tissue type from the binary metapeak vectors. We used five-fold cross-validation to estimate generalization accuracy, and inverse class-frequency weights to mitigate imbalance across tissue types. We modeled the same 19 tissue types as in our enrichment analysis, that is, those containing at least 10 samples. In L1-penalized regression, a parameter C controls a trade-off between prediction accuracy and model complexity, in terms of the number of metapeaks with non-zero weight. We varied C over values from 0.01 to 10, and observed the cross-validated accuracies shown in Figure 5A. Accuracy starts increasing significantly between C values of roughly 0.02 and 0.06, depending on the histone mark, and stabilizes when C is approximately 1. This saturation of performance occurred when the logistic model was selecting somewhere between 100 and 1000 metapeaks to make its predictions—far fewer than the total number of metapeaks we found to be significantly enriched in different tissue types. Notably, performance was strongly mark-dependent: H3K27ac achieved the highest plateau accuracy, followed by H3K4me1, H3K4me3, and H3K36me3, whereas H3K9me3 lagged substantially and required larger metapeak sets to approach its plateau. These trends indicate that activation-associated chromatin profiles more consistently encode tissue-discriminative regulatory programs than constitutive or broadly deployed repression-associated signal in this dataset.

**Fig. 5.**
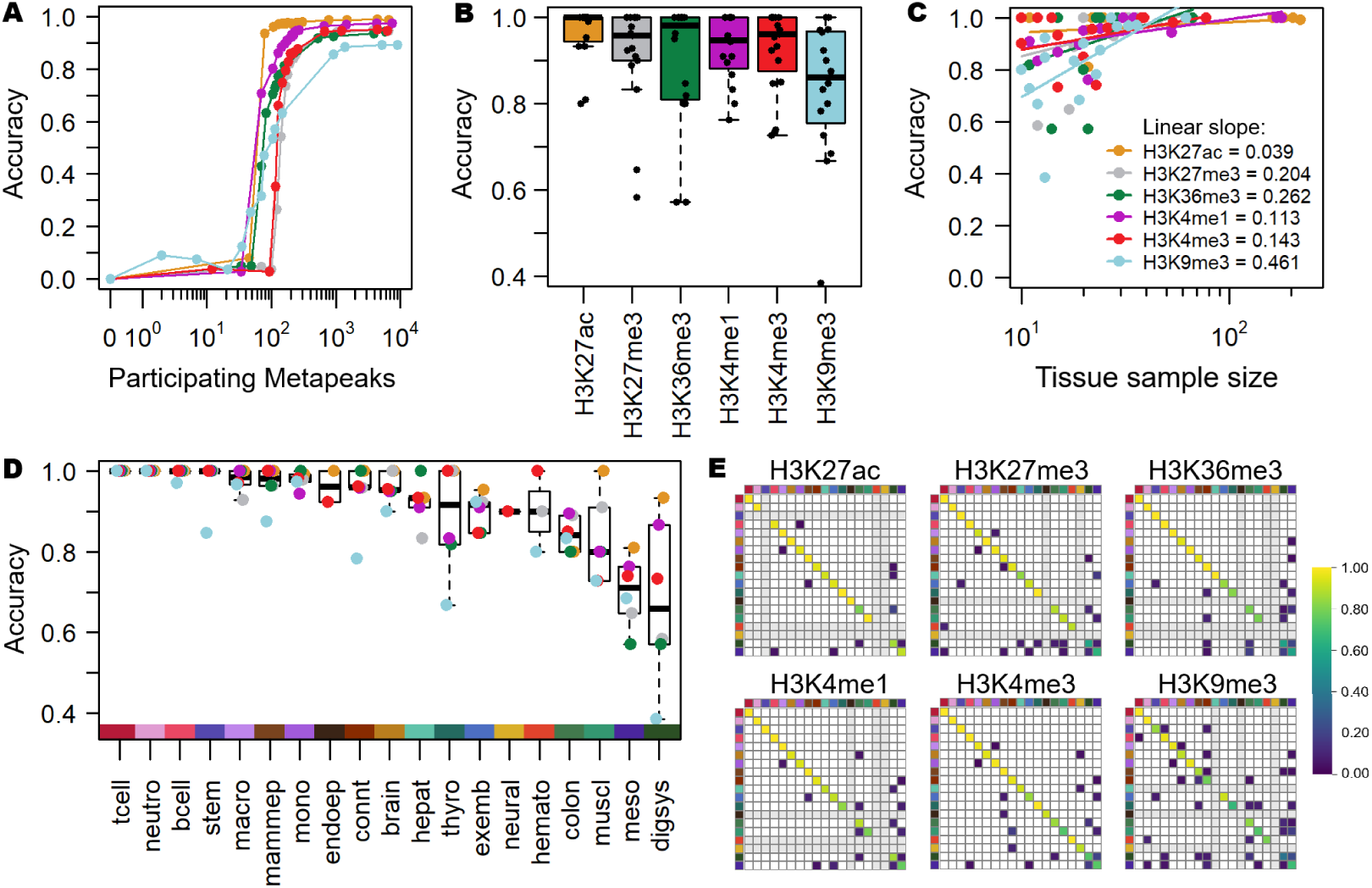
Metapeak-tissue links identified by machine learning accurately predict most cell/tissue types. A) Average prediction accuracy across all cell/tissue types for logistic regression with cross-validation (LR-CV) results for each histone mark, for model complexity C values ranging from 0.01 to 10. B) Box and whisker plots for cell/tissue type accuracies for each histone mark for LR-CV results based on a C value of 1. C) Dot plot with linear regression lines of LR-CV cell/tissue prediction results for each histone mark, showing the trend of accuracy with increasing sample size of the underlying cell/tissue types. D) LR-CV prediction accuracies grouped by cell/tissue type with box & whisker plots. Dots are colored by histone mark. Cell/tissue types are sorted by decreasing median accuracy. E) LR-CV confusion matrices shown as heatmaps. Zero values are white while large values follow a blue-green-yellow gradient. Cell/tissue types with *<* 10 samples were not tested, and are denoted by the off-white colored cells. Correct predictions appear along the diagonal lines of the confusion matrix. Tissues are denoted by the color sidebar.

Our hyperparameter optimization suggested that a model complexity penalty weight of *C* = 1 was where overall accuracy reached a plateau, offering near-best performance with the sparsest set of metapeaks. Figure 5B shows the accuracies on each tissue for each histone mark’s model. H3K27ac exhibited the strongest overall performance, with a median accuracy near 1.0 and the tightest inter-tissue spread, indicating robust generalization across diverse biological contexts. In contrast, H3K9me3 produced the lowest median accuracy of approximately 0.86 and broader dispersion. Despite these differences, most marks achieved high median accuracies (≥0.95 for all except H3K9me3), demonstrating that tissue identity is recoverable from metapeaks of different chromatin modalities.

To assess whether performance differences were driven in part by the number of samples available for each tissue type, we plotted cross-validated accuracies against the number of samples per histone mark-tissue type pair (Figure 5C). Most marks showed only weak-to-moderate dependence on sample size, with H3K27ac displaying minimal linear trend (slope ≈ 0.04), suggesting that its strong performance is not simply a function of larger class counts. In contrast, H3K9me3 demonstrated the strongest positive association with sample size (slope ≈ 0.46), consistent with greater sensitivity to limited training data and/or higher within-class variability for this repression-associated mark. Collectively, these relationships argue that mark-specific biological signal content, rather than sample size alone drives much of the performance, while also highlighting that H3K9me3 classifiers benefit disproportionately from increased sample size.

Performance varied strongly by tissue type (Fig. 5D), with several cell/tissue types being predicted accurately across essentially all marks. In particular, brain and multiple immune lineages (including T cells, B cells, macrophages, and neutrophils) were among the most consistently well-classified groups, indicating that they harbor distinctive metapeak patterns across both activation- and repression-associated landscapes. Conversely, certain categories—most prominently digestive system (digsys) and mesoderm-derived (meso) samples—were more frequently misclassified across marks, implying either closer regulatory similarity to other categories in the panel, higher heterogeneity within these groups, or incomplete separation at the granularity of our tissue labels.

Confusion matrices clarify the structure of the prediction errors and reveal biologically sensible failure modes (Figure 5E). Overall, H3K27ac yielded the cleanest diagonal (highest fidelity), including strong discrimination of closely related myeloid lineages. Macrophages were predicted with near-perfect accuracy, with only rare monocyte vs. macrophage confusion despite their very similar cell states (Geissmann et al, 2010). A representative, interpretable error was observed for colon under H3K27ac, where a minority of colon samples were miscalled as digestive system—an anatomically coherent relationship—while the reverse confusion was not prominent, suggesting asymmetric overlap between the colon-specific and broader digestive regulatory programs. Across marks, recurrent error hotspots again included digestive system and mesoderm-derived categories (final two rows and columns), reinforcing these as intrinsically challenging categories, at least under metapeak-based linear decision boundaries.

We wondered whether the metapeaks selected by machine learning were the same as those identified by our tissue-enrichment analysis, so we overlapped the two sets. Averaged across histone marks, 28% of metapeaks selected by machine learning were also enriched for a specific tissue, ranging from 46% of machine learning-selected H3K27ac metapeaks to 11% of H3K9me3 metapeaks. Conversely, therefore, over half of selected metapeaks were not enriched for a single tissue. This suggests that the machine learning was selecting metapeaks that might be enriched in multiple tissues. For instance, it might select a metapeak that is present in all blood-lineage cells, and rely on other metapeaks to discriminate between the blood cell types.

To connect predictive features to biological functions, for each cell/tissue type, we collected all positively-weighted metapeaks from any activation associated mark (H3K27ac, H3K4me1, H3K4me3, H3K36me3) and all negatively-weighted metapeaks for repression-associated marks (H3K27me3, H3K9me3). We then analyzed that set of regions using GREAT to test for enriched GO Biological Process terms. The full enrichment results are available in Supplementary File 5, but they largely mimicked the biologically-plausible results we obtained from our tissue enrichment analysis. For example, for brain, top terms emphasized nervous system biology (e.g., nervous system development, neurogenesis, neuron differentiation, learning/memory). For T cells, enriched terms centered on immune activation and adhesion programs (e.g., regulation of T cell activation, regulation of leukocyte cell–cell adhesion), consistent with lineage identity and signaling function. However, enrichment specificity varied across tissues: among immune subsets, neutrophils were the most readily distinguished by neutrophil-related terms in top-ranked results, while T cells and B cells shared multiple lymphocyte activation themes and were differentiated primarily by the presence/absence of a smaller number of lineage-leaning terms. Monocytes and macrophages showed less distinctive top-10 term sets in this analysis. Finally, not all tissues produced intuitively specific annotations, even when statistical enrichment was strong (e.g., hepatocyte-associated terms that were significant yet semantically broad), underscoring that GO term granularity and gene-set labeling can limit interpretability despite robust predictive signal. Still, taken together, these results show that logistic regression based on metapeak vectors yields high-accuracy tissue classification, especially for activation-associated marks, while using many fewer metapeaks than one-at-a-time enrichment analysis. Furthermore, the resulting signed feature sets often recover biologically sensible functional enrichments, while also delineating tissue categories and closely related lineages where additional modeling resolution or complementary annotations may be beneficial.

## 3 Discussion

In this study, we introduce FindMetapeaks, a scalable “peaks-of-peaks” framework for de novo discovery of recurrent histone-modification loci across thousands of uniformly processed ChIP-seq datasets spanning six major histone marks in the IHEC compendium. By treating high-confidence MACS2 peaks from each sample as the input “reads” for a second round of peak calling, we collapse billions of raw intervals into a compact, mark-specific atlas of metapeaks while retaining hallmark genomic properties of each mark and enabling downstream sample representation in a shared metapeak feature space.

A central result is that metapeaks preserve biologically meaningful structure at multiple resolutions: across all marks, the metapeak representation supports robust stratification of samples by histone-mark identity, while within each mark, cell/tissue identity emerges as the dominant axis of variation. Importantly, this tissue structure is not merely qualitative. Enrichment testing based on epi-allele frequency yields large catalogs of tissue-upregulated loci. A separate, machine learning-based approach independently recapitulates strong tissue-discriminative signal across marks. Together, these analyses argue that metapeaks provide a principled compression of large epigenomic repositories that retains interpretable regulatory content, rather than collapsing signal into an overly generic consensus.

One particularly useful interpretation of these results is that each tissue carries a reproducible “epigenetic fingerprint” in metapeak space, and that fingerprints differ both by cell/tissue type and by histone mark. In the EAF framework, tissue-upregulated metapeaks represent large-effect, tissue-skewed loci after controlling for ubiquitous signal via an explicit effect-size metric (dEAF). In the logistic regression framework, tissue-informative loci are identified by signed model weights that can be aggregated into tissue-specific metapeak sets and linked to gene programs. The convergence of these two perspectives—thresholded enrichment (Fig. 4) and discriminative modeling (Fig. 5)—supports the view that tissue identity is encoded by coherent, mark-aware regulatory architectures that can be summarized by a tractable set of genomic regions.

Our findings align with, and extend, prior large-scale efforts to define cross-sample regulatory loci (Feingold et al, 2004; Bernstein et al, 2005; Ernst et al, 2011; Kundaje et al, 2015; Meuleman et al, 2020). Conceptually, FindMetapeaks sits between simple union and intersection strategies. The union is often too permissive, while intersection is too stringent and can collapse the search space to a small set of near-ubiquitous regions with limited discriminatory utility. This intermediate position parallels prior work that clusters or consolidates regulatory intervals across many samples (e.g., the DNase hypersensitivity atlas of Meuleman et al (2020)) and is consistent with the broader trend in consortium-scale epigenomics toward building reusable coordinate systems for comparing heterogeneous experiments.

Our approach also contrasts with common uses of the term “metapeaks” in the literature, where it often refers to averaged signal profiles or heatmaps over peak sets (González et al, 2015; Keller et al, 2021; Savadel et al, 2021), rather than a de novo identification of recurrent loci across thousands of datasets. In addition, ENCODE-style replicate handling—intersection of replicates plus irreproducible discovery rate filtering—addresses a different problem (within-condition reproducibility) than ours (cross-condition recurrence and tissue informativeness). These strategies are therefore complementary rather than competing. Finally, the emerging “peak-calling-on-peaks” viewpoint has begun to appear in other large-scale contexts (Kudron et al, 2024), and the shared core idea strengthens the case that second-round peak calling can act as a general-purpose consolidation primitive when the number of datasets makes direct cross-sample integration unwieldy.

Several methodological choices and caveats deserve emphasis. First, we restricted meta-peak calling to the top 10,000 peaks per sample to reduce imbalance in peak counts across datasets, but this necessarily deprioritizes weaker yet potentially consistent loci that might recur broadly below the per-sample top-P threshold. Our software allows any choice of P, and also allows top-P selection to be turned off, so that all source-data peaks are used in metapeak calling. Second, most downstream analyses use a binary metapeak representation (peak present/absent in a metapeak) and do not incorporate quantitative peak strength, MACS2 pileup, or fold-enrichment information. Incorporating such quantitative signal could improve sensitivity and potentially sharpen tissue boundaries, particularly for marks or tissues that display heterogeneous signal (Hauduc et al, 2025). Third, we used narrow-peak calls throughout. Broad-peak calling might better summarize large areas of peak enrichment and further reduce the number of metapeaks and reduce redundancy.

Relatedly, there are plausible alternative representations we did not adopt. One could define a unified metapeak coordinate system across all marks by applying FindMetaPeaks to the peaks for all histone marks combined, although this would lose some of the region-specificity (e.g. promoter versus gene body) of the metapeaks. More radically, one could bypass peak calls entirely and call metapeaks directly from aggregated read pileups (e.g., bedGraph/bigWig summaries), potentially recovering loci missed by peak calling. However, this would require careful normalization across experiments and substantial computational resources, and would shift the method’s practical emphasis from portability to maximal sensitivity. It is also less clear how control signal would be accounted for in such a scenario.

Interpreting disease associations, as in our cancer versus healthy analysis, requires additional caution. While we observe cancer-associated metapeak signal that is broadly consistent with known biology in several contexts, pan-tissue cancer/healthy contrasts are intrinsically confounded by differential representation of different tissues, and even tissue-restricted contrasts may be influenced by tissue composition at the cellular level. This mixture caveat is especially salient in bulk epigenomics and raises the possibility that some significant differences might reflect altered underlying cell-type distributions rather than direct epigenetic reprogramming within a matched cell population. Analysis of single-cell epigenomic data could clarify this issue (Schwartzman and Tanay, 2015; Chen et al, 2025).

Looking forward, several directions are immediately compelling. Increasing sample coverage per tissue and per histone mark will improve both metapeak calling stability and tissue-level modeling, particularly for marks that show stronger dependence on sample size in classification settings. Integrating single-cell epigenomic assays (Schwartzman and Tanay, 2015) or spatially-resolved data (Deng et al, 2022) offers a direct route to disentangling tissue composition and cell state influences on recurrent epigenomic modifications. In this setting, the metapeak atlas could serve as a shared coordinate system for projecting single cells or spatial spots into a compact regulatory feature space, thereby enabling cleaner case–control comparisons, refined tumor substructure analyses, and more mechanistic hypotheses linking recurrent loci to cell-state transitions.

## 4 Methods

### Data

ChIP-seq peak data were downloaded from an sftp site shared internally among IHEC members, but the data is now also available from the IHEC Data Portal at https://epigenomesportal.ca/ihec/. Supplementary File 1 specifies precisely the datasets we used. For Figure 3D, gene, exon and intron definitions were obtained from Ensembl v94. Enhancer definitions used were the GeneHancer ‘double elite’ enhancers (Fishilevich et al, 2017). Repeat regions were obtained from the UCSC Genome Browser (Navarro Gonzalez et al, 2021). All data are in hg38 coordinates.

### Software

FindMetapeaks is implemented in Python and is available at https://github.com/rmbioinfo83/FindMetapeaks/. It depends on also having local installations of bedtools (https://bedtools.readthedocs.io/en/latest/) (Quinlan and Hall, 2010) and MACS2 (https://pypi.org/project/MACS2/) (Zhang et al, 2008).

### Analysis details

Views of peak pileups and called metapeaks in Figure 2 were generated in Integrated Genomics Viewer (IGV) version 2.19.4 (Robinson et al, 2011). Peak pileup tracks were generated as bedGraph files from bed files of metapeaks using the genomecov function. In IGV they were viewed with log-scaled y-axis and autoscaling. All other figure panels were generated using R or Python. tSNE embeddings (Figure 3E,F,H) were obtained using the tsne function in R, using default parameters. Heatmaps (Figures 3I, 4A-F) were hierarchically clustered using Euclidean distance with complete linkage. Logistic regression analyses were carried out in Python using sklearn’s LogisticRegression class with inverse tissue sample size weights and k=5-fold cross validation (Pedregosa et al, 2011).

## 6 Supplementary information

**Supplementary File 1: Supp1 IHEC Peak Sets.xlsx:** This spreadsheet lists all IHEC ChIP-seq peak sets used in our analysis, identified by file name as well as unique UUID. Also listed are the source tissue sample ID EpiRRID, histone mark that was assayed, cell/tissue type, and healthy/disease status.

**Supplementary File 2: Supp2 Metapeaks.xlsx:** This spreadsheet contains six sheets, one for each histone mark. Each sheet lists all metapeaks identified by FindMetapeaks by chromosome, start and end position in the first three columns. The remaining columns report on the ubiquitous, tissue-enriched, and machine learning-selected status of each metapeak.

**Supplementary File 3: Supp3 UbiquitousMetapeaks GREAT analyses.xlsx:** This spreadsheet contains six sheets, one for each histone mark. Each sheet lists the GREAT analysis results for the ubiquitous metapeaks, defined as having cross-tissue average epi-allele frequency of at least 0.9.

**Supplementary File 4: Supp4 TissueEnrichedMetapeaks GREAT analyses.xlsx:** This spreadsheet contains 18 sheets, one for each tissue type for which tissue-enriched metapeaks were identified. Each sheet lists the GREAT analysis results for the combined tissue-enriched metapeaks for the activation-associated marks (H3K27ac, H3K4me1, H3K4me3, H3K36me3).

**Supplementary File 5: Supp5 MLMetapeaks GREAT analyses.xlsx:** This spreadsheet contains 19 sheets, one for each tissue type studied in the machine learning analysis. Each sheet lists the GREAT analysis results for the positively-weighted activation-associated metapeaks and negatively-weighted repression-associated metapeaks identified by the machine learning.

**Supplementary Figure 1: FigS1 ROCCurve.pdf:** This figure displays a ROC analysis that quantifies the extent to which different metadata columns are clustered in the tSNE coordinates of Figure 3E,F.

