## Supplementary Figure S1 for "Histone Modification Metapeaks are Epigenetic Landmarks Predictive of Cell State"

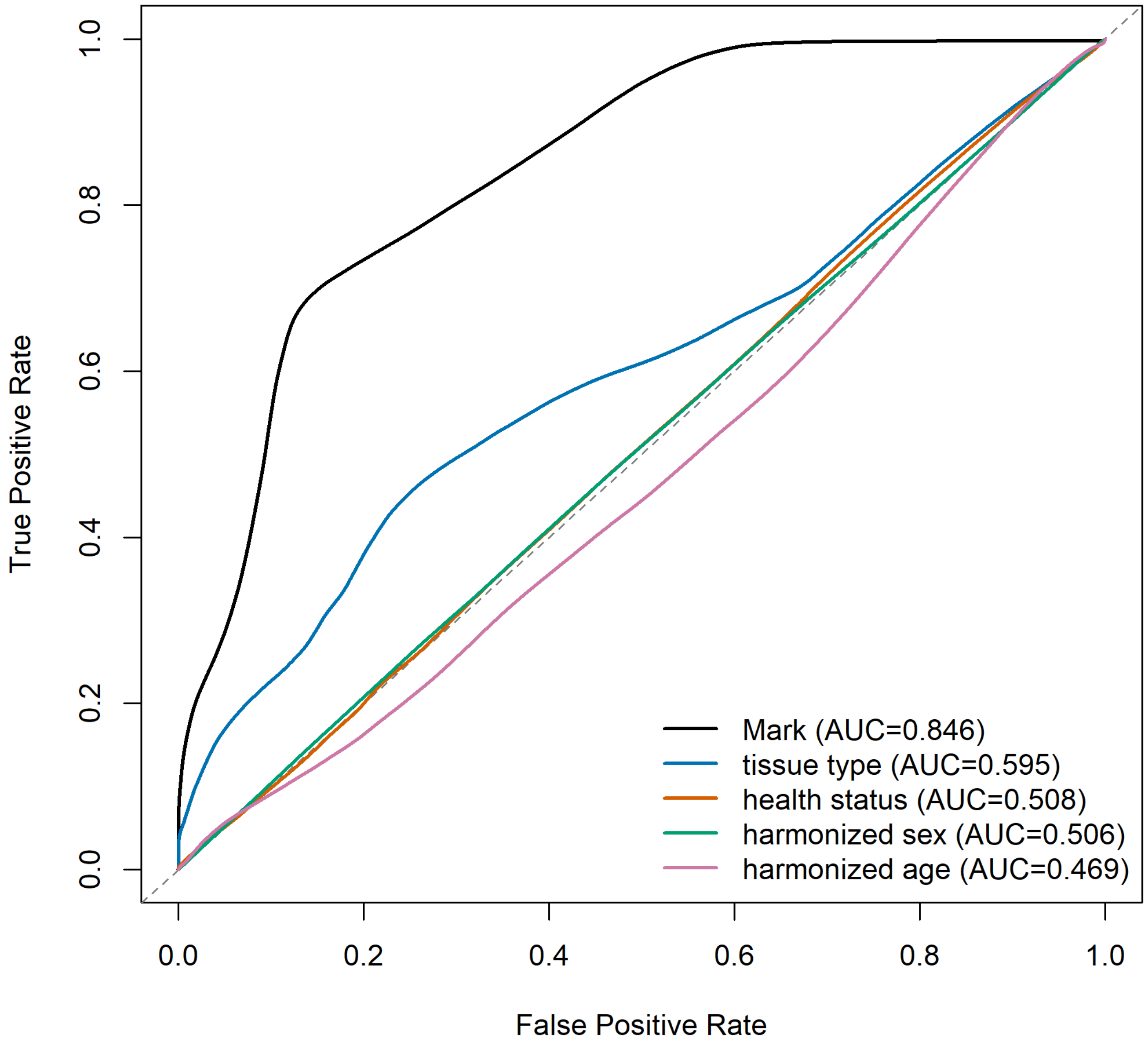


**Figure S1. ROC curves derived from pairwise sample correlations.** Merged metapeak regions shared across histone marks were used to construct a binary occupancy matrix for each experiment, where 1 indicates the presence of a peak within a metapeak region and 0 indicates its absence. Pairwise correlations were then calculated between all experiments using these binary vectors and ranked from strongest to weakest. ROC curves were generated to assess how well the ranked pairwise correlations recovered shared sample attributes, including histone mark, tissue type, health status, harmonized sex, and harmonized age.
